# Human milk extracellular vesicles target nodes in interconnected signalling pathways that enhance oral epithelial barrier function and dampen immune responses

**DOI:** 10.1101/2020.04.29.068841

**Authors:** Marijke I. Zonneveld, Martijn J.C. van Herwijnen, Marcela M. Fernandez-Gutierrez, Alberta Giovanazzi, Anne Marit de Groot, Marije Kleinjan, Toni M.M. van Capel, Alice J.A.M. Sijts, Leonie S. Taams, Johan Garssen, Esther C. de Jong, Michiel Kleerebezem, Esther N.M. Nolte-’t Hoen, Frank A. Redegeld, Marca H.M. Wauben

**Affiliations:** Department of Biomolecular Health Sciences, Faculty of Veterinary Medicine, Utrecht University, Utrecht, The Netherlands; Division of Pharmacology, Department of Pharmaceutical Sciences, Faculty of Science, Utrecht University, Utrecht, The Netherlands; Host-Microbe Interactomics Group, Department of Animal Sciences, Wageningen University, Wageningen, The Netherlands; Division of Immunology, Department of Infectious Diseases & Immunology, Faculty of Veterinary Medicine, Utrecht University, Utrecht, The Netherlands; Department of Experimental Immunology, Academic Medical Center, University of Amsterdam, Amsterdam Infection & Immunity Institute (AI&II), Amsterdam, The Netherlands Centre for inflammation, Amsterdam, The Netherlands; Centre for Inflammation Biology and Cancer Immunology, Dept Inflammation Biology, School of Immunology & Microbial Sciences, King’s College London, London, United Kingdom; Nutricia Research, Utrecht, The Netherlands

## Abstract

Maternal milk is nature’s first functional food. It plays a crucial role in the development of the infant’s gastrointestinal (GI) tract and the immune system. Extracellular vesicles (EVs) are a heterogeneous population of lipid bilayer enclosed vesicles released by cells for intercellular communication and are a component of milk. Recently, we discovered that human milk EVs contain a unique proteome compared to other milk components. Here, we show that physiological concentrations of milk EVs support epithelial barrier function by increasing cell migration via the p38 MAPK pathway. Additionally, milk EVs inhibit agonist-induced activation of endosomal Toll like receptors TLR3 and TLR9. Furthermore, milk EVs directly inhibit activation of CD4+ T cells by temporarily suppressing T cell activation without inducing tolerance. We show that milk EV proteins target key hotspots of signalling networks that can modulate cellular processes in various cell types of the GI tract.

---

In the neonate, the epithelial barrier of the GI tract needs to grow and mature, while the adaptive immune system is still developing^1^. This requires cellular regulation and education which, in part, is induced by components in mother’s milk. Although the importance of breastfeeding is widely recognized, the milk components and molecular processes involved in these developmental processes remain largely obscure^2^. This limited understanding is mainly due to the complexity of milk as it is composed of a wide range of bioactive macromolecular structures with partially overlapping physical properties^3^-^5^. Among these components are extracellular vesicles (EVs)^5-7^, which are nanosized particles released by cells. The release of EVs and the incorporation of molecular cargo into EVs is tightly regulated by the producing cell, resulting in a great heterogeneity of EVs specifically tailored for targeted intercellular communication^8,9^. Altogether this contributes to the versatile capacity of EVs to modulate cellular responses at multiple levels. The multi-faceted nature of EVs allows for the fine-tuning of complex signaling pathways in order to control the magnitude, kinetics and duration of cellular responses^10,11^. Previously, we have established an isolation procedure specifically designed for the reliable isolation of EVs from human milk^7^ and performed in-depth proteomics analysis of milk EVs^5^. We discovered several milk EV-associated proteins not identified before in human milk, that potentially contribute to both the development of the epithelial barrier and maintenance of immune homeostasis^5^. In this study, we explored the physiological role of milk EVs in these processes and linked their protein cargo to underlying signaling pathways.

Although often overlooked, the first interaction of milk components with the infants mucosa occurs in the oral cavity, a site that needs to maintain an intact epithelial barrier and contains mucosa-associated lymphoid tissue (MALT)^12,13^. To determine whether milk EVs play a role in maintenance of the physical epithelial integrity, we performed a gap closure assay with gingival epithelial cells in the presence of milk EVs. Milk EVs significantly increased the re-epithelialization rate of epithelial cells to almost the same level as the positive control TGF-α, while EV-depleted milk supernatant did not (Fig. 1A and 1B, Supplementary File 1). Re-epithelialization can occur through p38 MAPK-dependent migration^14,15^, either or not combined with cell proliferation through MEK-ERK signaling^16^. To assess which milk EV proteins could be involved in the enhanced re-epithelialization and which pathways were likely targeted, we performed enrichment, network- and functional annotation analysis on the milk EV proteome^5^. Using enrichment analysis we first identified 33 significantly enriched GO-terms involved in cell cycle and migration to which collectively 159 proteins were associated (Supplementary File 2). Interestingly, network analysis revealed that 134 of these 159 proteins can form protein-protein interactions (Supplementary Fig. 1A), indicating that the combined EV cargo can be delivered as protein networks. Finally, functional annotation analysis was performed to link each protein from the identified protein networks to relevant signaling cascades involved in reepithelization, while also visualizing the expected mode of action of the milk EV protein (Supplementary Fig. 1B). These analysis resulted in a model that shows that milk EV proteins can interact at multiple levels in signaling cascades and unveils several ‘hot spots’ where milk EV proteins formed nodes with a high number of potential interactions. Several of these nodes are formed around key cellular proteins that are involved in either inhibition of cell cycle (and proliferation), or stimulation of migration via p38 MAPK (Supplementary Fig. 1B and summarized in Fig. 1C). To investigate whether milk EVs primarily affected re-epithelialization by stimulating migration, cells were cultured in the presence of pharmacological inhibitors of p38 MAPK, or MEK1/2 as a control for proliferation. Inhibition of p38 MAPK abolished the increased re-epithelialization rate induced by EVs (Fig. 1D), while MEK1/2 inhibition showed only a minor reduction of EV-mediated re-epithelialization (Fig. 1E). Interestingly, others have shown a role for milk EVs of rat, porcine and bovine origin in modulating intestinal epithelial cells^17-19^, suggesting that supporting epithelial integrity could be a central and evolutionary conserved function of maternal milk EVs. This idea is further supported by the presence of conserved milk EV miRNA cargo with regulatory functions that are shared between mammalian species, which could aid in this process^20^. Taken together, we demonstrate that human milk EVs enhance gingival epithelial cell migration via p38 MAPK. Milk EV proteins likely form clusters that have the potential to target proliferation and migration signaling cascades at multiple levels, thereby modulating the balance between key signaling nodes and tailoring cellular responses.

**Fig. 1:**
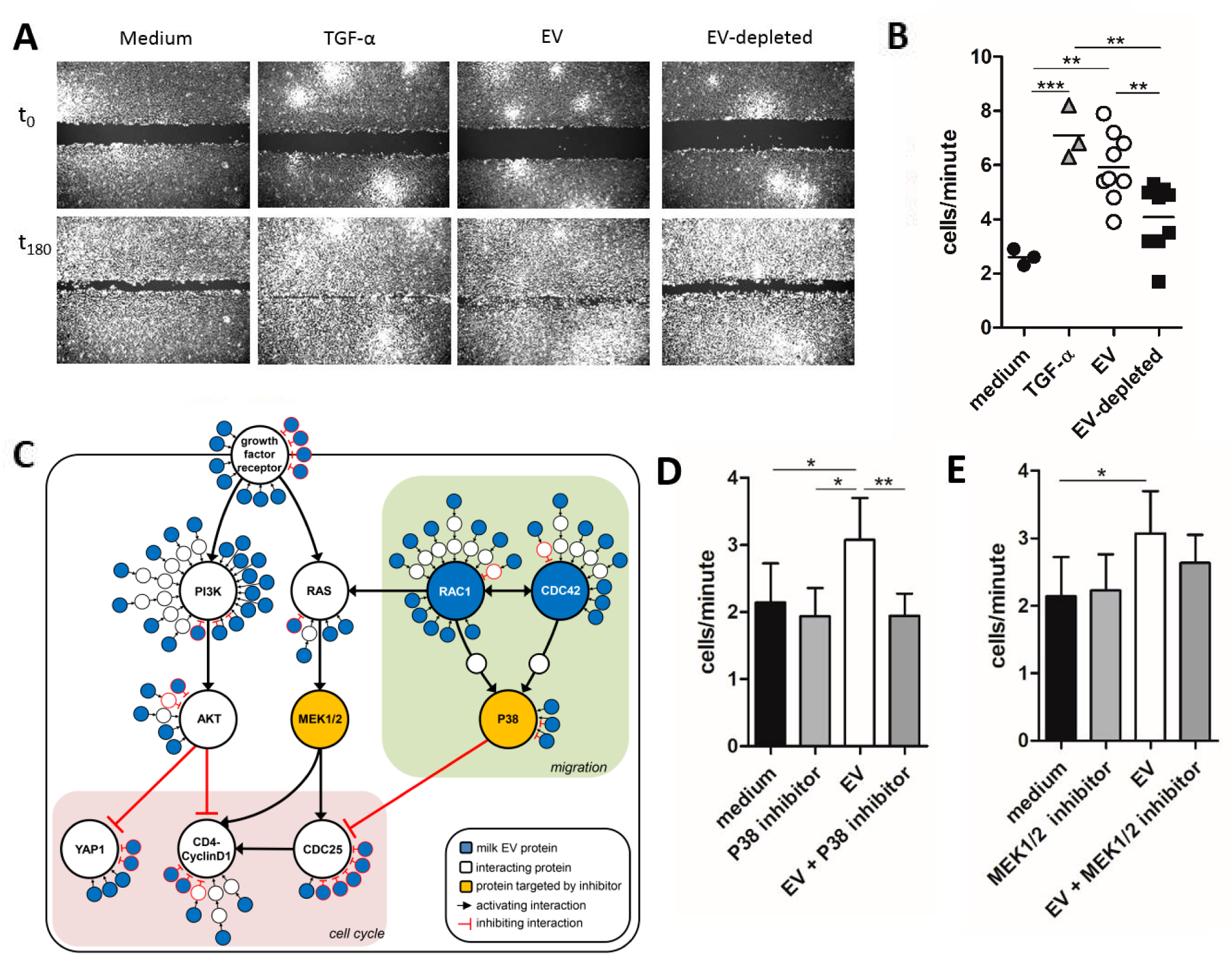
Human milk EVs enhance the formation of the epithelial barrier via p38 MAPK. A-D) A gap was made in a confluent monolayer of Ca9-22 gingival epithelial cells and re-epithelialization was investigated in the presence of milk EVs, or milk donor-matched EV-depleted procedural and biological control, or TGF-α as a positive control. Cell migration was tracked in triplicate conditions by live cell imaging. Images were acquired every 20 minutes for 5 hours, or until the gap in the TGF-α condition was closed. A) Representative microscopic images of Ca9-22 cells at the start of experiment (t0) and after 180 minutes (t180) incubation with indicated conditions. Images are representative of 2 experiments with 3 milk donors per experiment. B) Quantification of Ca9-22 cells migrating into the gap, expressed in cells per minute during the linear growth phase. Each well was plotted as a single point and data are representative of 2 experiments with 3 milk donors in each experiment. C) Schematic representation of key signaling pathways involved in gap closure. From the full annotation analysis (Supplementary file 2 and Supplementary Fig. 1), key signaling pathways were selected that could be involved in gap closure. Several nodes with high interactions (≥ 6 interactions) were observed. These hotspots were involved in either i) inhibition of cell cycle (e.g. YAP1, CDK4-CyclinD1 inhibition via the PI3K-AKT pathway, or CDC25 inhibition via p39 MAPK pathway), ii) activation of cell cycle (e.g. MEK1/2 via RAS), or iii) stimulation of migration (e.g. the p38 MAPK pathway via RAC1 and CDC42). Proteins present in milk EVs are depicted in blue, while interacting cellular proteins are shown in white. Key proteins involved in cell cycle or migration that were inhibited in D and E are highlighted in yellow. Type of interactions between proteins are shown as activating or inhibiting. D-E) Quantification of Ca9-22 cells migrating into the gap in the presence of P38 MAPK inhibitor (D) or MEK1/2 inhibitor (E). Data is representative of 2 independent experiments performed in triplicate with 2 different milk donors per experiment. Bars represent mean ± SD. Significance was calculated by one-way ANOVA with * p < 0.05; ** p < 0.01 and *** p < 0.001.

Epithelial cells lining the oral cavity also act as sentinels by scouting their environment via innate immune receptors. The activation of such receptors, e.g. Toll-like receptors (TLRs), needs to be critically regulated in order to enable the mucosal system to appropriately defend against pathogens, but tolerate commensals^12,21,22^. We therefore investigated whether milk EVs could be involved in TLR regulation.

Using TLR reporter cell lines, we observed that milk EVs stimulated both TLR2 and TLR4 to some extent (Fig. 2A and 2B), while TLR3 and TLR9 were only triggered by their respective agonist (Fig. 2C and 2D). Although it has previously been shown that EVs from other sources can stimulate TLR2^23^, TLR3^24^, TLR4^23^ and TLR9^25^, we show that milk EVs do not activate TLR3 and TLR9. Next, we determined whether milk EVs could regulate agonist-induced TLR activation. Both TLR2 and TLR4 reporter lines showed equal stimulation in the presence of agonist regardless of the presence of milk EVs (Fig. 2A and 2B). In contrast, milk EVs significantly dampened the agonist-induced response of endosomal TLR3 and TLR9, while EV-depleted controls had no or reduced effects (Fig. 2C and 2D). To assess the inhibitory effect of milk EVs on endosomal TLR activation in a more physiological setting, we evaluated their effect on gingival epithelial cells. As these epithelial cells do not express TLR9, we assessed inhibition of TLR3 activation and found that also in these cells milk EVs reduced TLR3 activation, as measured by gene transcription of the pro-inflammatory cytokines *IL6* and *CXCL8* (Fig. 2E). The expression profile of other TLR-related genes showed that in the presence of milk EVs, agonist-induced gene upregulation could be fully inhibited (*LTA, REL, MAP2K4* and *IL6)*, or partially inhibited (*MAPK8, PELI1, FOS, JUN*, and *CXCL8*) (Fig. 2F and Supplementary Fig. 2 for complete gene array). In contrast, milk EVs enhanced the agonist-induced upregulation of *EIF2AK2*, as well as that of *SIGIRR and SARM1*, both negative regulators of TLR signaling^26,27^. The enhancing effect of milk EVs was also observed for several genes that were downregulated by the agonist (*IRAK4, TLR6, PRKRA, PPARA* and *TLR3*). Finally, opposite effects of agonist-induced activation were observed in the absence or presence of milk EVs (*TRAF6* and *TIRAP)* (Fig. 2F). Collectively, the gene expression data indicate that milk EVs do not passively prevent TLR triggering by sequestering TLR ligands, but that they modulate TLR responses in an active manner. To assess which milk EV proteins could be involved in TLR modulation, we performed enrichment, network-, and functional annotation analysis and identified 26 significantly enriched GO-terms with 130 EV proteins associated to TLR signaling, of which 110 proteins formed protein-protein networks (Supplementary File 2 and Supplementary Fig. 3A). From this analysis we extracted the key TLR signaling pathways targeted by milk EVs, leading to cytokine production or regulation of TLR trafficking and processing (Supplementary Fig. 3B and Fig. 2G). Based on this model, the TLR2 and TLR4 activating effect of milk EVs in the absence of TLR agonists (Fig. 2A and 2B) can be explained by the presence of several TLR2 and TLR4-stimulating proteins associated to milk EVs. As the signaling cascades for both plasma membrane and endosomal TLRs converge, no distinctive regulatory mechanism could be identified in our model that explains the inhibition of endosomal TLRs signaling.

**Fig. 2:**
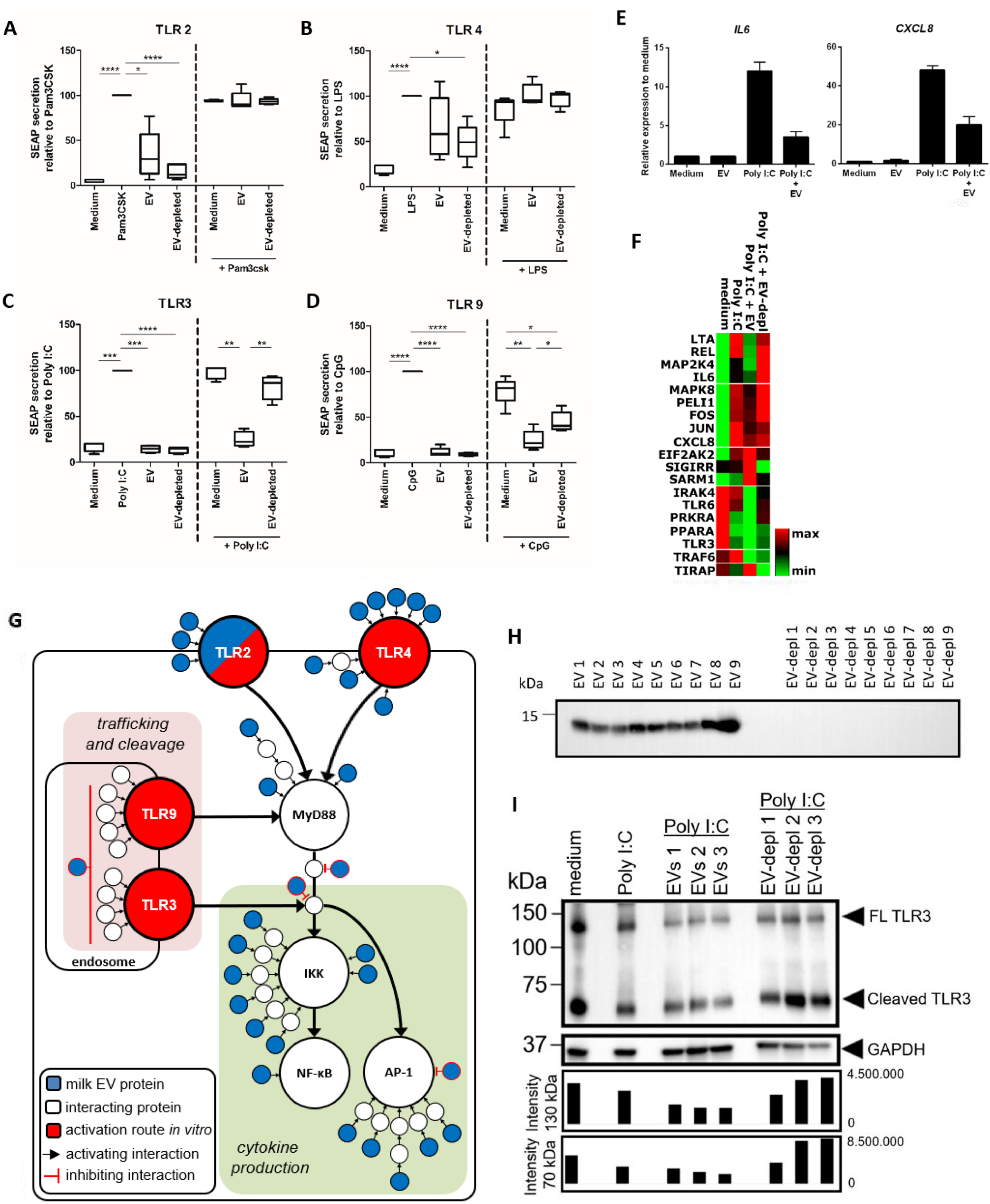
Human milk EVs inhibit agonist-induced activation of endosomal TLR3 and TLR9, but not of cell surface TLR2 and TLR4. A-D) Secretion of SEAP reporter protein was determined for TLR reporter cell lines cultured in indicated conditions either or not in response to agonist (Pam3CSK for TLR2, LPS for TLR4, Poly I:C for TLR3, and CpG ODN2006 for TRL9), with the TLR-specific agonist set to 100%. Box and whisker plots contain data of 3 independent experiments performed in triplicate with a total of 4 different milk donors. E) Relative gene expression of IL6 and CXCL8 in Ca9-22 cells cultured for 5 hours with medium, EV, Poly I:C or Poly I:C + EV. Delta Ct-values to ACTB were calculated and expressed relative to medium controls. EVs from 2 different milk donors were used and PCR reaction was performed twice. Results are summarized in bar graphs as mean ± SD. F) Heatmap representing the cellular gene expression profile of Ca9-22 cells cultured for 4 hours in the 4 different conditions (medium, agonist, EV + agonist, EV-depleted + agonist) in a gradient running from minimal gene expression (green) to maximal expression (red) for each gene analyzed (see Supplementary Fig. 2 for complete dataset). Data is derived from 1 experiment, with 1 milk donor. G) Schematic representation of key signaling pathways involved in TLR signaling of the individual TLRs tested which are shown in one Figure. TLR2, TLR4 and TLR9 signal via MyD88, while TLR3 signals via TRIF (not shown) to NF-kB (via IKK) and/or AP-1 leading to cytokine production. TLR2 and TLR4 are surface receptors, while TLR3 and TLR9 are sorted into the endosome where they are cleaved which enhances their signaling. Proteins from milk EVs are depicted in blue, while interacting cellular proteins are shown in white. Type of interactions between proteins are either shown as activating or inhibiting. In every assay, the individual TLR was activated via its specific ligand, which is shown in red. Note that TLR2 is present in milk EVs as well as in the TLR2 reporter cells. H) Western blot analysis for the presence of Cystatin-B (CSTB; expected size 11 kDa; exposure time 19 seconds) in purified milk EVs or in EV-depleted control. Data are from 9 individual milk donors. I) Western blot analysis for the presence of TLR3 on whole cell lysate of Ca-922 cells cultured in medium alone or stimulated with poly I:C with or without milk EVs or EV-depleted control. Full length (FL) TLR3 (expected size 130 kDa) and cleaved TLR3 (expected size 70 kDa) are visible. GAPDH (expected size 36 kDa) was used as a loading control and applied to normalize the intensity of the full length band and cleaved band (shown as volume intensity below the blot) in order to quantitively compare signals. A total of 3 different milk donors were tested. Significance was calculated by one-way ANOVA with * p < 0.05; ** p < 0.01; *** p < 0.001 and **** p < 0.0001.

However, in contrast to TLR2 and TLR4, endosomal TLRs require specific trafficking and processing which involves transportation into the endosome followed by proteolysis via cathepsins, which enhances their signaling^28-30^. From our functional annotation analysis, we identified the cathepsin inhibitor cystatin-B (CSTB)^31^, and verified its presence in milk EVs using immunoblotting (Fig. 2H). Although CSTB has not been reported to be involved in the modulation of endosomal TLR cleavage, we explored whether milk EVs could affect the unprocessed full length and the cleaved form of TLR3 after agonist-induced activation of epithelial cells. Indeed, we observed less cleaved TLR3, however the abundance of full length TLR3 was also lower (Fig. 2I). This suggests that milk EVs can modulate expression, or might interfere with TLR3 synthesis, for which the inhibition of *TLR3* gene expression is indicative (Fig. 2F). Taken together, our data show that milk EVs selectively inhibit agonist-induced endosomal TLR signaling, regulate downstream gene expression and reduce cellular full length and cleaved TLR3 expression.

The selective regulation of key hotspots in TLR signaling could support the colonization of the mucosa in the newborn, as it has been shown that differential NF-κB activation is essential for immune homeostasis and tolerance to commensal bacteria^22,32^.

In contrast to innate immune cells, cells from the adaptive immune system are educated during postnatal development, whereby CD4+ T cells located in the MALT differentiate into T helper (Th) and regulatory subsets^33,34^. Since epithelial cells can endocytose milk EVs^35^ and selectively transport macromolecular structures from the apical to basolateral compartments via transcytosis^36^, it is assumed that milk EVs can reach cells underlying the epithelial barrier. Therefore, we evaluated the effect of milk EVs on T cell activation and differentiation. When CD4+ T cells were activated in the presence of milk EVs, they were halted in their proliferation (Fig. 3A and 3B). In order to determine whether milk EVs only interfered in proliferation and not differentiation, we monitored the ratio of naïve CD45RA+ to memory CD45RO+ T cells^34,37^. As expected, activation of T cells skewed the T cell population towards CD45RO at the expense of CD45RA (Fig. 3C and 3D). In contrast, addition of milk EVs during T cell activation prevented this switch and T cells retained CD45RA expression (Fig. 3C and 3D). The inhibitory effect of milk EVs was not merely via blocking proliferation of memory T cells (CD45RO), as proliferation of highly purified naïve CD4+CD45RA+ T cells was also inhibited (Supplementary Fig. 4). This indicates that milk EVs inhibit activation of naïve T cells and their transition to a memory phenotype. Further evidence for impaired T cell activation in response to milk EVs was found by the inhibited release of a wide range of different Th subset-associated cytokines, e.g. IFN-γ (Th1), IL-5 (Th2), IL-9 (Th2, Th9 and Th17), IL-10 (Th2 and regulatory T cell: Treg), IL-13 (Th2), IL-17A and IL17F (Th17), and IL-22 (Th1, Th17) (Fig. 3E). Suppression of proliferation and cytokine production could be indicative for Treg induction^34^, as Treg inhibit T cell activation and it has been previously reported that a crude preparation human milk EVs induced expression of the Treg-associated transcription factor FoxP3^6^. Yet, after exposure of activated T cells with milk EVs, we did not observe the induction of the typical Treg phenotype with high expression of CD25 and FoxP3 in the absence of CD127 (Supplementary Fig. 5). Additionally, we functionally tested the inhibitory capacity of T cells exposed to milk EVs. For this, CD4+ T cells were first stimulated in the presence of milk EVs, recovered and cultured with fresh CD4+ T-responder cells. As expected, in this suppression assay the milk EV-primed T cells were not suppressive (Fig. 3F). To asses which pathways might be targeted by milk-EV proteins in the observed inhibition of T cell activation, we used enrichment analysis and identified a total of 48 significantly enriched GO-terms containing 165 proteins, of which 137 proteins were part of protein-protein interaction networks (Supplementary File 2 and Supplementary Fig. 6A).

**Fig. 3:**
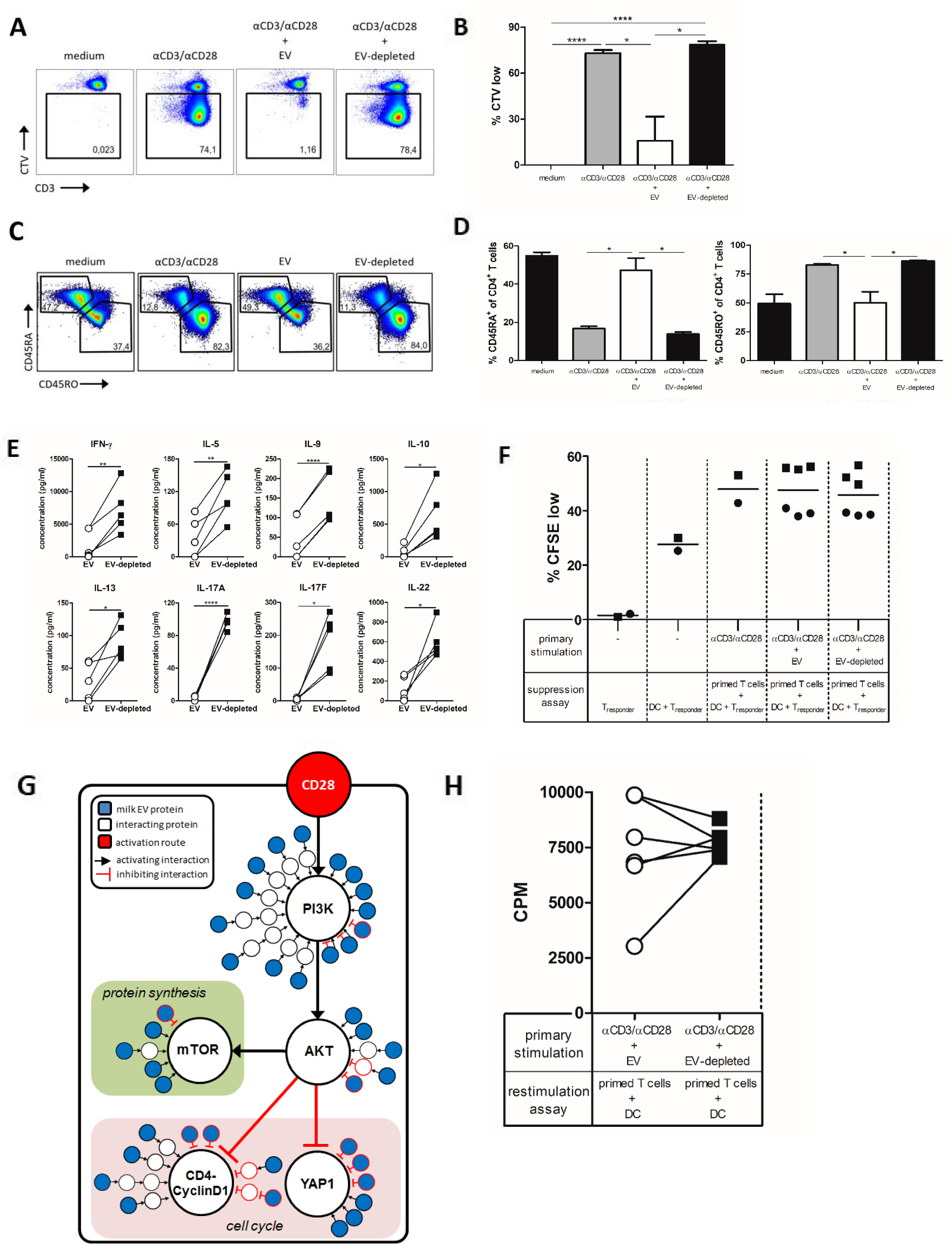
Human milk EVs transiently inhibit CD4+ T cell activation and retain cells in a naïve phenotype. A-D) Purified CD4+ T cells were labeled with CellTrace Violet (CTV) and incubated with medium, αCD3 and αCD28 (αCD3/αCD28), or αCD3/αCD28 in the presence of EV or EV-depleted control for 6 days. A) Representative dot plots of CTV dilution in response to the indicated conditions. Percentage CTV^low^ cells (gate) is expressed as a fraction of total CD4+ T cells. B) Quantification of the percentage CTV^low^ CD4+ T cells following incubation with the indicated conditions. Bars summarize mean ± SD of 2 independent experiments using different T cell donors (n=2) and different milk donors (n= 2 and 3). C) Representative dot plots of CD4+ T cells stained for CD45RA and CD45RO after culture in the indicated conditions controls for 6 days. Percentage of CD45RA+/CD45RO- and CD45RO+/CD45RA-cells in gates are expressed as a fraction of total CD4+ T cells. D) Quantification of the percentage CD45RA+ and CD45RO+ cells of total CD4 T cells following incubation with the indicated conditions. Bars summarize mean ± SD of 2 independent experiments using different T cell donors (n=2) and different milk donors (n = 2 and 3). E) Cytokine profiles of stimulated CD4+ T cells incubated with EV or EV-deleted controls. The following values were obtained for αCD3/αCD28 stimulated controls (not shown in the bar graph), provided as mean ± SD: IFN-γ: 4514 ± 1757 pg/ml, IL-5: 89 ± 30 pg/ml, IL-9: 123 ± 37 pg/ml, IL-10: 330 ± 243 pg/ml, IL-13: 99 ± 22 pg/mL, IL-17A: 13 ± 8 pg/ml (compared to 97 ± 8 pg/ml with EV-depleted), IL-17F: 68 ± 47 pg/ml (compared to 180 ± 76 pg/ml with EV-depleted), IL-22: 346 ± 59 pg/ml. Data is shown for day 6 as highest secretion was seen for αCD3/αCD28 controls on this day. Graphs summarize results of 2 independent different T cell donors (n=2) and different milk donors (n = 2 and 3). F) Suppression assay performed with CFSE-labeled CD4+ Tresponder cells incubated with allogeneic monocyte-derived DC and autologous CD4+ T cells that received primary stimulation with αCD3/αCD28 or with αCD3/αCD28 in the presence of EVs or the EV-depleted control. After priming, the cells were washed, irradiated and added to an allogeneic mixed lymphocyte reaction (MLR). Plotted are single points for the average of technical triplicates, shown as a percentage of total Tresponder cells. Data is representative of 2 independent experiments (with 2 individual donors) using 3 different milk donors. G) Schematic representation of key signaling pathways involved in regulating T cell activation: downstream signaling of CD28 with inhibition of in cell cycle (e.g. YAP1 and CDK4-CyclinD1 via PI3K-AKT) and activation of mTOR (via PI3K-AKT). Proteins from milk EVs are depicted in blue, while interacting cellular proteins are shown in white. Type of interactions between proteins are either shown as activating or inhibiting. CD28 is highlighted in red, as this receptor was stimulated. H) Restimulation assay with purified CD45RA+ CD4+ T cells (n=2) that received a primary stimulation with αCD3/αCD28 in the presence of EVs or EV-depleted control (n=3). Cells were subsequently restimulated with allogeneic monocyte-derived DC in the absence of EV and EV-depleted controls. Restimulation was performed in triplicate and the average values plotted as a single point. Proliferation was measured by ^3^H-thymidine incorporation and data is expressed as counts per minute (CPM). Significance was calculated with one-way ANOVA (B, D and F), or paired t-test (E and H) and significance defined as * p < 0.05; ** p < 0.01 and **** p < 0.0001.

Using functional annotation analysis we could link the majority of these interacting proteins to relevant signaling cascades downstream of CD28, resulting in the inhibition of cell cycle and the stimulation of mTOR (Supplementary Fig. 6B and Fig. 3G). This model clarifies the inhibition of proliferation and cytokine production by milk EVs, as the inhibition of downstream CD28 signaling can result in the retained naïve CD4+ T cell phenotype^38^. Furthermore, the absence of Treg induction by milk EVs becomes evident, as mTOR and PI3K inhibit FoxP3 expression in T cells^39^. These findings suggest that milk EVs directly inhibit CD4+ T cell activation, without T cell tolerance or Treg induction. To test this, we activated naïve CD4+ T cells in the presence of milk EVs, recovered the cells and restimulated them with allogeneic DC. Indeed, T cells that were primed in the presence of milk EVs, proliferated after restimulation, to the same extent as restimulated T cells primed with EV-depleted control (Fig. 3H). This demonstrates that the presence of milk EVs causes a transient inhibition of T cell activation which is reversible. Based on our findings we propose that milk EVs create a temporary increased threshold for CD4+ T cell activation. Raising this threshold helps the infant to cope with the high antigenic load to which it is exposed after birth creating tolerogenic conditions that are optimal for development^40,41^.

Although human milk is known to aid the postnatal development and maturation of the GI tract, and maintenance of immune homeostasis^1,42-44^, the exact effects on the oral mucosa has remained understudied. Here we demonstrate that purified EVs from human milk can promote gingival cell re-epithelialization, modulate epithelial endosomal TLR responses and transiently control T cell activation (Supplementary Fig. 7). As EVs are heterogeneous in composition, the net-effect on the target cell will be determined by the collective EV cargo delivered and relevant pathways targeted. Therefore, EV-induced effects result from the concerted action of numerous biomolecules delivered, rather than from a single regulatory molecule. This is underscored by our models derived from the functional annotation analysis of the milk EV proteome, in which potential interactions of protein cargo occurs throughout the signaling cascades and where nodes of high interaction affect key signaling proteins. Whereas some ‘regulatory hotspots’ like PI3K, are prominent in all three biological processes studied, other unique hotspots were identified in each biological assay. In that respect we postulate that for the biological function of EVs, heterogeneity of the EV pools ensures fine-tuning with respect to the specific cellular response. Future studies are needed to determine which EV subsets interact with specific recipient cells as it is likely that EV subsets will contain specific cargos that are geared towards distinct functions in target cells, and that their composition is controlled by environmental cues from the mother^45^.

In summary, EVs are multifactorial and bioactive components of human milk that can interact with various cell types found in the oral mucosa, creating a window of opportunity for regulated development of the epithelial barrier and the innate and adaptive immune system of the newborn.

## Methods

### Human milk collection

Human milk was collected as previously described^7^. Briefly, fresh and mature milk samples were collected by 16 healthy mothers, with a mean age of 33 ± 2.4 years who were at a lactational stage of 3 to 15 months (with an average of 7.2 ± 3 months). The milk was prevented from cooling down and EV isolation started within 30 minutes after collection. Informed consent was signed by all donors and this study was approved by the local ethics committee.

### Human milk EV isolation and EV-depleted milk control isolation

Isolation of milk EVs was performed as previously described with some modifications for functional analysis^7^. The workflow is described in Supplementary Fig. 8A. Whole milk was centrifuged at 22°C for 10 minutes at 3,000 g (Beckman Coulter Allegra X-12R, Fullerton, CA, USA). After removal of the cream layer the harvested milk supernatant was centrifuged at 3,000 g again at 22°C and stored at −80°C until further processing. Thawed supernatant was centrifuged in polyallomer SW40 tubes (Beckman Coulter) at 5,000 g for 30 minutes at 4°C and subsequently at 10,000 g (Beckman Coulter Optima L-90K with a SW40Ti rotor). Next, 6.5 ml aliquots of the 10,000 g supernatant were loaded on top of a 60% – 10% iodixanol gradient (Optiprep™, Progen Biotechnik GmbH, Heidelberg, Germany) in a SW40 tube. Gradients were ultracentrifuged at 192,000 g (Beckman Coulter Optima L-90K with a SW40Ti rotor) for 15-18 hrs. Following this centrifugation, the resulting EV-depleted milk supernatant (6.5 ml) left on top of the gradient was collected and loaded onto a new iodixanol gradient and ultracentrifuged once more and further processed identically to milk EV samples, to obtain a donor-matched procedural milk control. The EV and EV-depleted samples from the iodixanol gradients were collected in fractions of 500 µl and densities of 1.06-1.19 g/ml were pooled, as these had a high expression of the EV-associated marker CD9 in Western blot for the EV sample (Supplementary Fig. 8B). Iodixanol was removed from samples by size exclusion gel filtration using a 20 ml column (Bio-Rad Laboratories, Hercules, CA, USA) packed with 15 ml Sephadex g100 (Sigma-Aldrich, St. Louis, MO, USA) and elutriating 24 fractions of 1 ml with phenol red free RPMI 1640 or DMEM medium (Gibco™, Invitrogen, Carlsbad, CA, USA). Eluates 3 – 9 contained EVs as these had a high protein concentration (data not shown) and a high expression of the EV-associated marker CD63 (Supplementary Fig. 8C). The EV-containing eluates were pooled and supplemented with 10% heat-inactivated fetal calf serum (FCS; Sigma-Aldrich), or 0.1% BSA (Sigma-Aldrich), 2mM ultraglutamine (BioWhittaker, Lonza, Switzerland), and 100 IU/ml penicillin and 100 mg/ml streptomycin (Gibco). Samples were frozen at −80°C until use. The isolated milk EVs were characterized by Western blot for EV-associated markers CD9, CD63, Flotillin-1 and HSP70, as well as the non-EV marker Lactoferrin (Supplementary Fig. 8D). Concentrations of EVs used for *in vitro* assays were within or below the physiological range, as the starting volume of the 10,000 g milk supernatant (6.5 ml) from which the milk EVs were isolated is in the range of the pooled eluates (7 ml). We have submitted all relevant data of our experiments to the EV-TRACK knowledgebase (EV-TRACK ID: EV200007)^46^.

### Re-epithelialization assay

Ca9-22 (JCRB0625) gingival epithelial cells (JCRB Cell Bank, Osaka, JP) were cultured in DMEM containing Glutamax (Gibco) and supplemented with 10% FCS (Gibco), 100 U/ml penicillin and 100 μg/ml streptomycin (Sigma-Aldrich). The re-epithelialization assay was performed as previously described^47^. For this, cells were seeded at 3.5 × 10^4^ cells/well in 96-well flat bottom tissue culture treated plates (BD Falcon™, Corning, NY, USA) and left to reattach and form a confluent monolayer overnight. Cells were starved in FCS-free DMEM for 2 hours prior to experiments and labeled during the last 20 minutes of starvation with 2 µM CellTracker™ Red CMTPX and 2 µg/ml Hoechst 33342 (both from Molecular Probes, Eugene, OR, USA). A gap was made in the cell monolayers of every well using an HTS Scratcher (Peira, Antwerpen, BE). Cells were washed twice with PBS (Gibco) and cultured in FCS-free DMEM as non-treated control, 4 ng/ml human transforming growth factor α (TGFα; R&D Systems, MN, USA) as a positive control, or with 100 μl milk EVs, or 100 μl EV-depleted control. For inhibition of migration via p38 MAPK or proliferation via MEK1/2, 10 μM of p38 inhibitor (SB203580) or MEK1/2 inhibitor (U0126) was used (Cell Signaling Technology, Danvers, MA, USA). Cells were monitored by live cell imaging using the BD Pathway 855 Bioimaging System (BD Biosciences). Images of the same frame of each well were acquired every 20 minutes for 5 hours or until the gaps in the positive controls were closed. Image segmentation and data analysis were carried out using CellProfiler 2.1.1 (https://www.cellprofiler.org/) and FCS Express 4 Plus (De Novo Software, Glendale, CA, USA). A nonlinear least squares regression was used to fit the modified Gompertz function^47^ through the re-epithelialization measurements obtained per well. The repair rate parameter derived from the model was used to calculate the average re-epithelialization rate (cells/min) from technical triplicates.

### TLR reporter assay

HEK-Blue™-hTLR2, HEK-Blue™-hTLR3, HEK-Blue™-hTLR4, and HEK-Blue™-hTLR9 reporter cell lines (Invivogen, Toulouse, FR) were cultured in DMEM medium containing Glutamax (Gibco), supplemented with 8.5% heat-inactivated FCS (Bodinco), 50 units/mL penicillin (Sigma-Aldrich), 50 µg/mL streptomycin (Sigma-Aldrich) and 100 µg/mL Normocin (Invivogen). Additionally, the HEK-Blue hTLR9 culture medium was supplemented with 10 μg/mL blasticidin (Invivogen) and 100 μg/mL zeocin (Invivogen), the HEK-Blue hTLR3 culture medium with 30 µg/mL blasticidin and 100 µg/mL zeocin and the HEK-Blue hTLR4 and – hTLR2 culture medium with 1x HEk-Blue Selection (Invivogen). For experiments, cells were cultured in 96-well flat bottom tissue culture treated plates with 90 µl milk EVs or EV-depleted with a final volume/well 110 µl. Agonists used were 100 ng/mL Pam3CSK, 5 μg/ml Poly I:C, 10 ng/mL LPS-EK, or 1 μg/ml CpG ODN2006 (all from Invivogen). Cells were cultured for 16 hours after which supernatant was harvested and SEAP reporter protein secretion was determined using QUANTI-Blue™ detection medium (Invivogen) and measuring absorption at 650 nm on a 550 Microplate reader (Bio-Rad) as per manufacturer’s instructions.

### TLR stimulation of Ca9-22 cells

Ca9-22 cells were seeded at 3.5 × 10^4^ cells/well in 96-well flat bottom plates (Corning) with DMEM medium containing Glutamax (Gibco) and supplemented with 10% FCS (Gibco) (Sigma-Aldrich), 100 U/ml penicillin and 100 μg/ml streptomycin (Sigma-Aldrich). The next day, the media was replaced by FCS-free medium containing 0.1% BSA (Sigma-Aldrich) and cells were stimulated with 100 μl milk EVs or EV-depleted controls in the presence or absence of 5 μg/ml Poly I:C (Invivogen). After 4 or 5 hours, cells were washed with PBS (Gibco), harvested and stored at −80°C in RLT buffer with β-mercaptoethanol from the RNeasy Mini prep Kit (Qiagen, Hilden, Germany) according to manufacturer’s instructions until cDNA synthesis for RT^2^ Profiler PCR array. For TLR3 and GAPDH Western blot, cells were stored in RIPA buffer (40mM Tris pH 8.0, 150mM NaCl, 1% Triton X-100, 0.5% Na-deoxycholate, 0.1% SDS composition) buffer until use.

### SDS page and Western blotting

For characterization of samples during EV isolation, 100 µl aliquots were collected from iodixanol gradient Optiprep fractions, or eluates from the size exclusion gel filtration column. Isolated milk EVs, EV-depleted sample, or fractions were pelleted by centrifugation for 65 minutes at 100,000 g (in a Beckman Coulter Optima Max-XP with a TLA-55 rotor) in polyallomer microcentrifuge tubes (Beckman). Pelleted fractions were resuspended in sample buffer (62.5 mM Tris pH 6.8, 2% SDS, 10% Glycerol), while pelleted EVs or EV-depleted sample was resuspended in RIPA buffer (40mM Tris pH 8.0, 150mM NaCl, 1% Triton X-100, 0.5% Na-deoxycholate, 0.1% SDS). Whole cell lysates from Ca9-22 cells were prepared from 5.2*10^5^ cells in 30 µl in RIPA buffer. Protein content of whole cell lysate was measured with the BCA protein assay (Pierce™, Thermo Scientific, Landsmeer, Netherlands) to quantify the protein content of the samples in order to equalize for input material for SDS page. Samples were heated at 95°C for 3 minutes and run on a 8 – 16% or 4 – 20% TGX-Criterion gel (Bio-Rad). The separated proteins were transferred to PVDF membranes and blocked in PBS containing 0.2% fish skin gelatin (Sigma-Aldrich) and 0.1% Tween-20. Proteins were detected by immunoblotting using mouse anti-human CD9 (clone HI9a, BioLegend, dilution 1:1.000), mouse anti-human CD63 (clone TS63, Abcam, dilution 1:1.000), mouse anti-human flotillin-1 (clone 18, BD Biosciences, dilution 1:5.00, sample reduced with β-mercapthoethanol) and mouse anti-human HSC70/HSP70 (clone N27F3-4, ENZO dilution 1:1.000, sample reduced with β-mercapthoethanol) and rabbit anti-human lactoferrin (polyclonal, Abcam, dilution 1:5.000), rabbit anti-human TLR3 (D10F10, Cell Signaling TECHNOLOGY, dilution 1.1000, sample reduced with β-mercapthoethanol), mouse anti-human GAPDH (mAbcam 4984, Abcam, dilution 1:1.000), mouse anti-human CSTB (clone 225228, R&D Systems, dilution 1:5.00, sample reduced with β-mercapthoethanol). Goat anti-mouse-HRP (Jackson Immuno Research, Suffolk, UK; dilution 1:10.000), or goat anti-rabbit-HRP (DAKO, dilution 1:1.000) was used as secondary antibody. HRP conjugated antibodies were detected using SuperSignal West Dura Chemiluminescent Substrate (Thermo Scientific and ChemiDoc XRS and Image Lab 5.1 (Bio-Rad).

### RT-qPCR and RT^2^ Profiler PCR array

Total RNA was extracted from Ca9-22 cells after 4 or 5 hours of culture using the RNeasy Mini prep Kit and cDNA was prepared using the RT^2^ First Strand Kit (Qiagen) or the qScript™ cDNA Synthesis Kit (Quanta Biosciences, MA, USA) according to the manufacturer’s instructions. Real-time quantitative PCR (RT-qPCR) was performed on a Rotor-Gene Q2plex real-time cycler (Qiagen). For relative gene expressions of *IL6* and *CXCL8*, delta Ct-values were log-transformed with *GAPDH* and *ACTB* as internal controls. For gene expression profiling, cDNA was added to the Human Toll-like receptor signaling pathway RT^2^ Profiler PCR array (Qiagen) and run on an iCycler MyiQ (Bio-Rad). The relative expression levels of each gene were normalized to the expression level of 5 reference genes included in the array (*ACTB, B2M, GAPDH, HPRT1 and RPLP0*). Delta Ct-values were log-transformed and analyzed by the web-based software GeneGlobe Data Analysis Center (Qiagen). Genes for which gene expression levels had a Ct>35 in all test conditions were excluded from analysis.

### CD4+ T cell isolation

Human peripheral blood mononuclear cells (PBMC) were isolated from Buffy coats by Lymphoprep™ density gradient centrifugation (Axis-shield, Dundee, United Kingdom and Nycomed, Zurich, Switzerland). PBMC were cultured in RPMI 1640 supplemented with Glutamax and sodium pyruvate (Gibco), 2.5% FCS and 100 IU/ml penicillin and 100 mg/ml streptomycin (Gibco). CD4+ T cells were isolated from PBMC using CD4+ T cell isolation kit (Miltenyi Biotec, Bergisch Gladbach, Germany) according to manufacturer’s instructions (average purity of 95%) and were used directly in experiments or fractionated into CD45RA+ and CD45RO+ subsets using anti-CD45RA-PE (UCHL-1, Dako) and anti-PE magnetic beads (Miltenyi Biotec) resulting in >97% pure T cell subpopulations.

### T cell stimulation

PBMCs were seeded at 4×10^6^ cells/ml in 12 wells plates (Corning) coated with 1.5 µg/ml αCD3 (CLB-T3/4.E, 1XE from Sanquin, Amsterdam, The Netherlands) and cultured in RPMI 1640 medium (Gibco) with 10% FCS (Sigma-Aldrich), 750 μl milk EVs or EV-depleted controls in a total volume of 1 ml and cultured for 6 days. Purified CD4+ T cells were seeded at 0.5×10^6^ cells/ml in 48 wells plates (Corning) coated with 0.5 – 1.5 µg/ml αCD3 (CLB-T3/4.E, 1XE from Sanquin, Amsterdam, The Netherlands) and cultured with 70 ng/ml – 1 µg/ml soluble αCD28 (CLB-CD28/1, 15E8 from Sanquin), RPMI 1640 medium (Gibco) with 10% FCS (Sigma-Aldrich), 750 μl milk EVs or EV-depleted controls in a total volume of 1 ml and cultured for the indicated amount of time.

### Multiplex cytokine analysis

Supernatants of stimulated T cells were harvested on day 6 of culture and analyzed using the LEGENDplex™ human Thelper 1/2/9/17 multiplex kit (BioLegend). Beads were acquired on a BD Canto II (BD Bioscience) and analyzed with LEGENDplex™ V7.0.

### T cell suppression assay

T cell suppression assay was performed as described previously^48^. In brief, CD4^+^CD45RA^+^ T cells were activated for 5-6 days with 1.5 µg/ml plate bound αCD3 (Sanquin) and 1 µg/ml soluble αCD28 (Sanquin) in medium, or in the presence of EV or EV-depleted controls. T cells were subsequently harvested, irradiated (3,000 rad), washed and counted. Cells were replated at a 2:1 ratio with 0.5 µM CFSE (Sigma-Aldrich) labeled autologous CD4^+^CD45RO^+^ responder cells in the presence of monocyte-derived allogeneic DC. Cells were incubated for 5-6 days and CFSE dilution of T cells was determined on a BD Canto II (BD Bioscience) and analyzed by FlowJo (v10.1: FlowJo, Ashland, OR, USA).

### Restimulation assay

10×10^6^ cells/ml CD4^+^CD45RA^+^ T cells were cultured for 3 days in the presence of 1.5 µg/ml plate bound αCD3 (CLB-T3/4.E, 1XE Sanquin) and 1 µg/ml soluble αCD28 (CLB-CD28/1, 15E8 Sanquin) in the presence or absence of milk EVs or EV depleted control. T cells were subsequently harvested, washed, counted and reseeded at a 4:1 ratio with allogeneic monocyte-derived DC generated as previously described^49^ in a mixed lymphocyte response. Cells were incubated for 3 days total and to determine T cell proliferation 11 KBQ/well ^3^H-thymidine (Radiochemical Center, Amersham, Little Chalfont, UK) was added. The incorporated ^3^H-thymidine was measured after 16 h by liquid scintillation spectroscopy.

### Flow cytometry

Proliferation of purified CD4+ T cells was assessed by labeling cells with 2 μM CellTrace Violet (Invitrogen) or 0.5 µM CFSE (Sigma-Aldrich) prior to culture. Following culture, cells were harvested and stained with fluorescent conjugated antibodies. Antibodies used in this study: CD3-PE-Cy7 (UCHT1, dilution 1:100), CD4-PerCP-Cy5.5 (OKT4, dilution 1:50), CD25-Alexa488 (BC96, dilution 1:20), CD45RA-PE (HI100, dilution 1:50), and CD45RO-APC-Cy7 (UCHL1, dilution 1:50), CD127-PE-Cy7 (A019D5, dilution 1:20) (all from BioLegend). FoxP3-APC (eBioscience, San Diego, CA, USA, 236A/E7, dilution 1:25). Intracellular staining for Treg phenotype were done using the Anti-human Foxp3 Staining Set (eBioscience) according to the manufacturers protocol. Cells were measured on a BD Canto II (BD Bioscience), and analyzed by FlowJo (v10.1 FlowJo, Ashland, OR, USA).

### Functional enrichment, network- and annotation analysis

Previously, we unraveled the common milk-EV proteome^5^, in which 367 proteins were identified in all 7 human milk EV samples tested. These proteins were loaded into the STRING (version 10.5) protein-protein interaction network analysis tool^50^, of which 362 proteins matched the STRING database (Supplementary File 2). Enrichment analysis for biological process on these proteins identified n=784 significantly enriched GO-terms, of which several linked to the observed *in vitro* effects of milk EVs. GO-terms were selected that linked to cell cycle (proliferation) and regulation of migration (n=33), or regulation of TLR signaling (n=26), or regulation of T cell activation (n=48) (Supplementary File 2 for data and complete workflow). As enrichment analysis alone does not provide information on the molecular action of a proteins, or its position in a signaling cascade, we next performed a comprehensive functional annotation analysis. For that purpose, signaling pathways underlying the observed *in vitro* effects were constructed using Uniprot entries and the associated sources from Uniprot itself for each individual selected milk EV protein from the enrichment analysis. This pathway analysis was supplemented with a general literature search on both the individual milk EV proteins, as well as the signaling pathways associated to the *in vitro* observations in order to prevent biased results solely from the enrichment data. Annotation analysis on the selected milk EV proteins was done to determine their interaction with other proteins (either given a common gene name, or a synonym when widely used in literature) and the type of interaction (activating or inhibiting). For each milk EV protein it was decided whether the protein itself, or its cellular interaction partner was relevant to signaling pathways involved in the particular *in vitro* experiments. Although we could link many milk EV proteins to underlying signaling cascades, yet for some proteins there was insufficient experimental evidence to do so (Supplementary File 2). In the end, all relevant signaling cascades from the full functional annotation analysis are depicted in supplementary figures (Supplementary Fig. 1B for proliferation and migration; Supplementary Fig. 3B for TLR signaling; ; Supplementary Fig. 6B for T cell activation), in which ‘hot spots’ were defined as a node in the signaling cascade where a cellular protein would have at least 6 potential interaction partners. Key pathways from the full annotation analysis are shown as a summary in the main figures whereas irrelevant signaling cascades are omitted for clarity. Additionally, protein-protein network analysis on the identified milk EV proteins that linked to the selected GO-terms was performed in STRING (minimum required interaction score set to high confidence 0.700), followed by k-means clustering in order to construct clusters of interaction proteins within the network.

### Statistical analysis

Data were analyzed by paired t-test or one-way ANOVA with Tukey’s multiple comparison test in GraphPad Prism Software V5.01. GO-analysis in String was tested with a Fisher’s exact test followed by a correction for multiple testing. Significance was defined as *p < 0.05; **p < 0.01; ***p < 0.001 and **** p < 0.0001.

## Supplementary Figures

**Supplementary Fig. 1:**
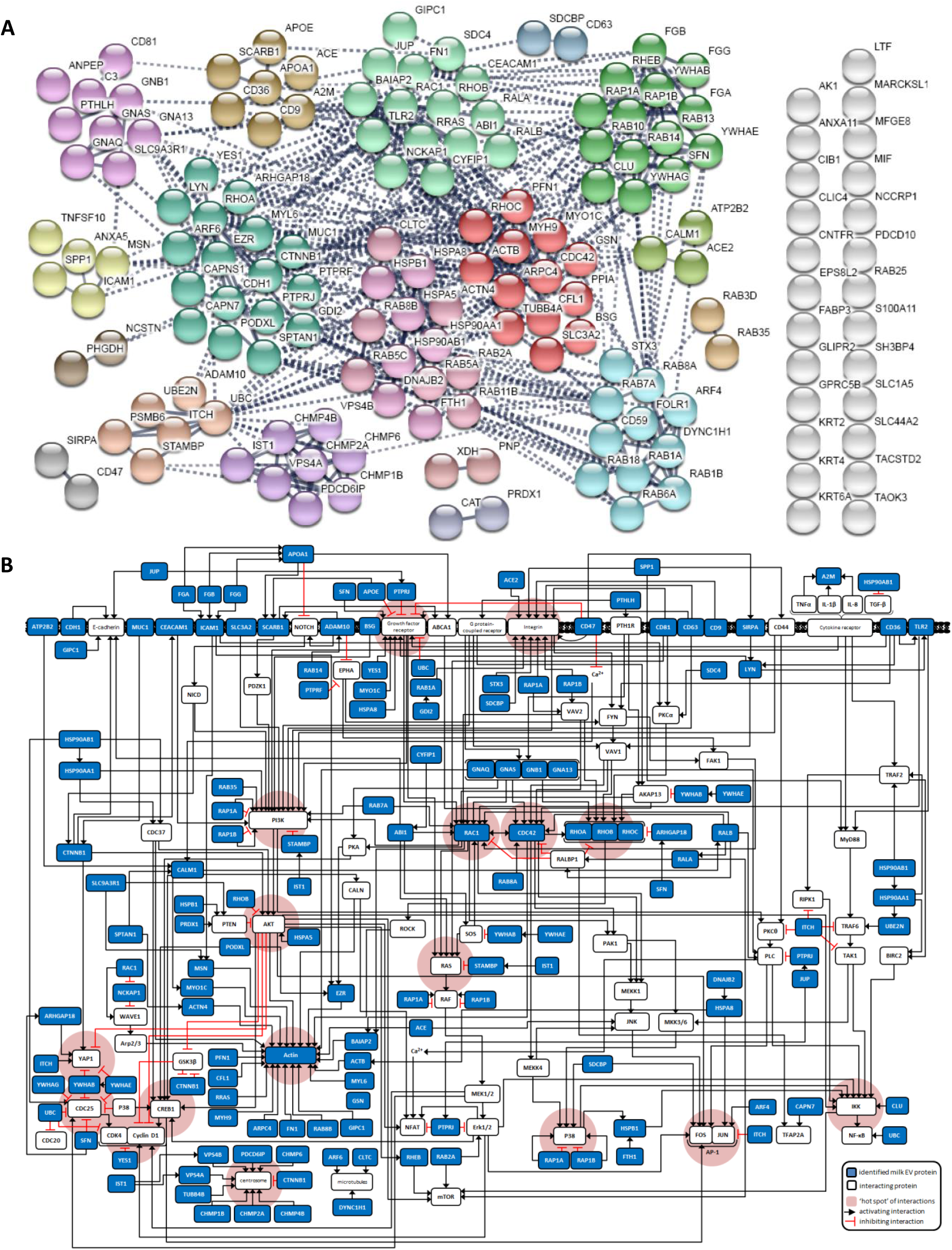
Enrichment and protein-protein network analysis allowed complete integration of human milk EV proteins into signaling pathways associated to cell cycle (proliferation) and migration and revealed interactions at multiple levels in the signaling cascades. A) Protein-protein interaction analysis on the identified milk EV proteins that linked to the selected GO-terms was performed (minimum required interaction score set to high confidence 0.700), followed by k-means clustering (showing 18 clusters) in order to visualize the most likely occurring clusters within the network. A total of 134 proteins formed protein-protein interactions (which is 84.3%) and 25 proteins had no interaction with any other milk EV protein (which is 15.7%). Only those proteins that were part of an interaction network were further investigated for validated links to proliferation or migration signaling cascades. B) Functional annotation analysis of milk EV protein clusters that link to cell cycle (proliferation) and migration. The interaction of selected milk EV proteins (in blue) and cellular proteins (in white; either shown with their common gene name, or a synonym when widely used in literature) and the type of interaction (activating or inhibiting) within relevant signaling pathways are shown. If a protein has interactions with ≥ 6 other proteins, this node is shown in red as a ‘hotspot’. Although milk EV proteins were selected via relevant GO-terms, some proteins could not be linked to the specific signaling cascades and are not shown in B (see Supplementary File 2 for a full overview of the analysis).

**Supplementary Fig. 2:**
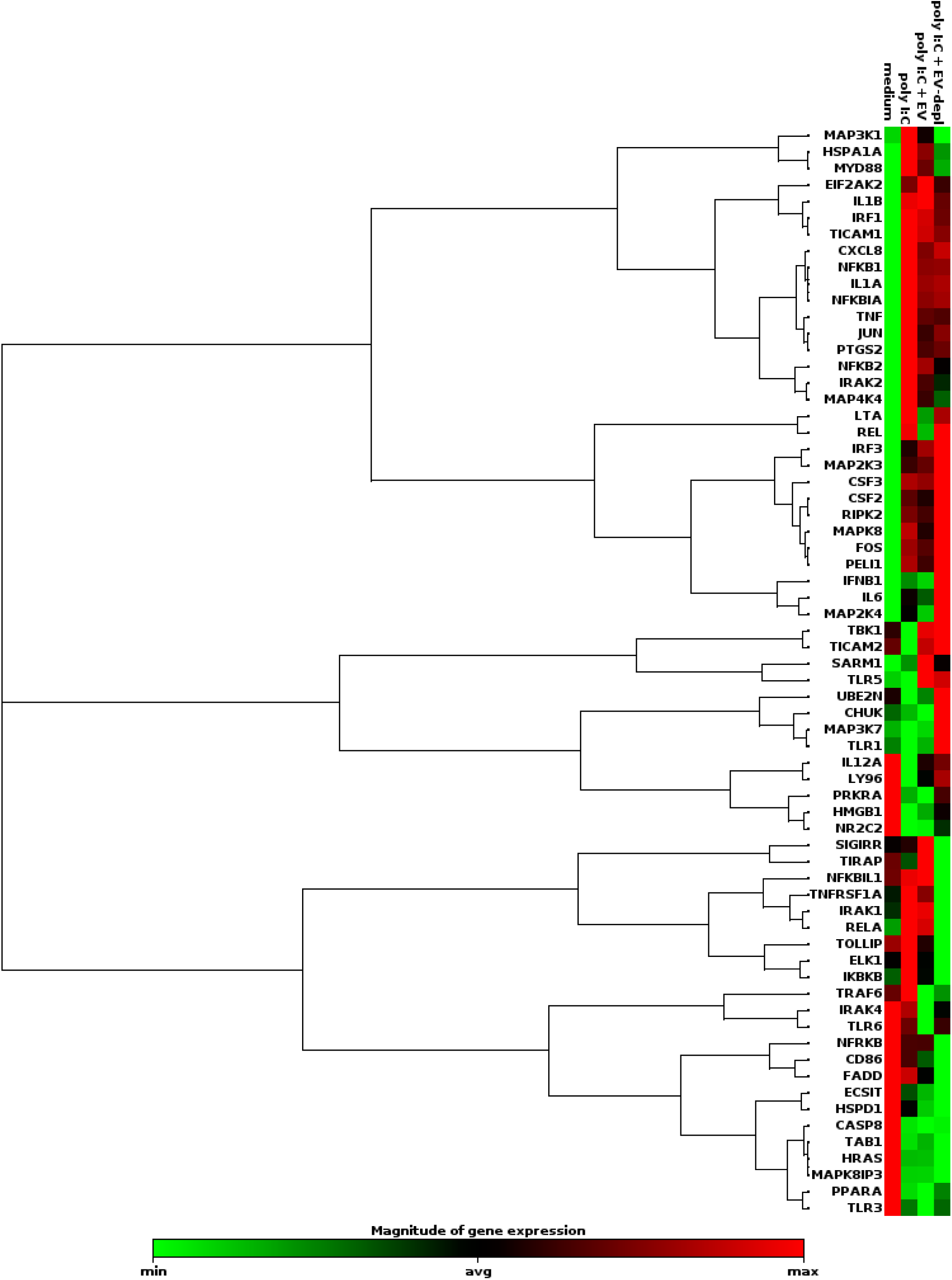
Complete heatmap of cellular gene expression profile from gingival epithelial cells exposed to human milk EVs. Complete dataset from gene array in Fig. 2F, which shows the non-supervised hierarchical clustering of all culture conditions to display a heat map with dendrograms indicating co-regulated genes across groups or individual samples. The heat map of gene expression is shown as a gradient running from minimal gene expression (green) to maximal expression (red) for each gene analyzed. Genes for which gene expression levels had a Ct>35 in all test conditions were excluded from the analysis and household genes are not shown.

**Supplementary Fig. 3:**
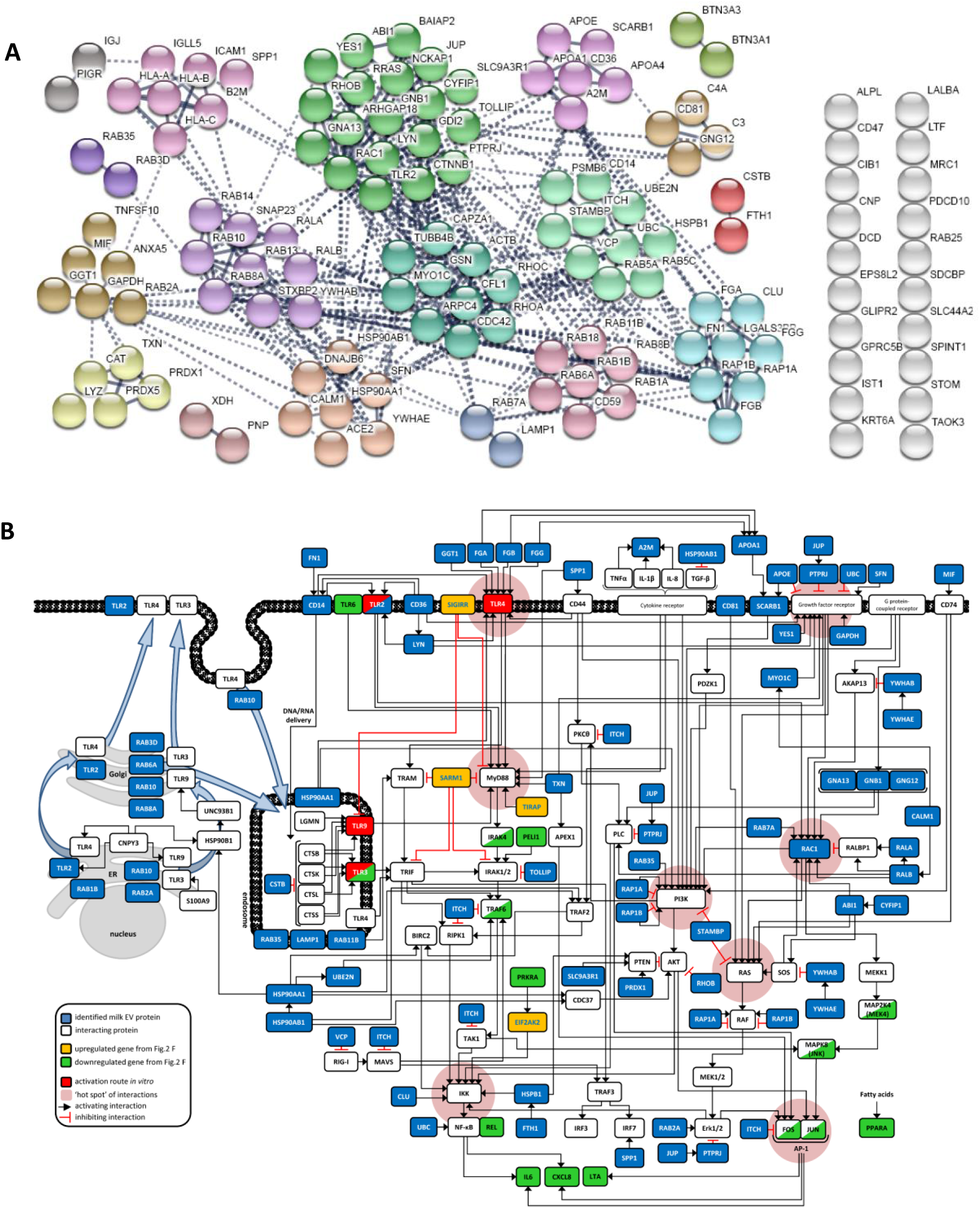
Enrichment and protein-protein network analysis allowed complete integration of human milk EV proteins into signaling pathways associated to TLR signaling and revealed interactions at multiple levels in the signaling cascades. A) Protein-protein interaction analysis on the identified milk EV proteins that linked to the selected GO-terms was performed (minimum required interaction score set to high confidence 0.700), followed by k-means clustering (showing 18 clusters) in order to visualize the most likely occurring clusters within the network. A total of 110 proteins formed protein-protein interactions (which is 84.6%) and 20 proteins had no interaction with any other milk EV protein (which is 15.4%). Only those proteins that were part of an interaction network were further investigated for validated links to TLR signaling cascades. B) Functional annotation analysis of milk EV protein clusters that link to TLR signaling. The interaction of selected milk EV proteins (in blue) and cellular proteins (in white; either shown with their common gene name, or a synonym when widely used in literature) and the type of interaction (activating or inhibiting) within relevant signaling pathways are shown. Furthermore, we included in the model the TLR-associated genes (Fig. 2F) that were differentially expressed after incubation of the epithelial cells with TLR3 agonist in presence of milk EVs which resulted in either upregulated or downregulated gene expression compared to medium, agonist, or EV-depleted control. If a protein has interactions with ≥ 6 other proteins, this node is shown in red as a ‘hotspot’. Although milk EV proteins were selected via relevant GO-terms, some proteins could not be linked to the specific signaling cascades and are not shown in B (see Supplementary File 2 for a full overview of the analysis).

**Supplementary Fig. 4:**
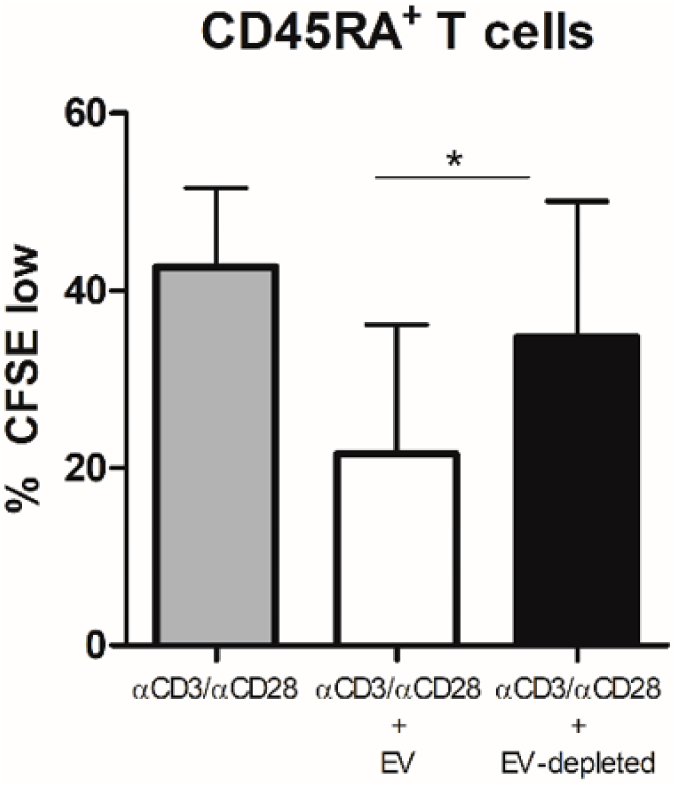
Human milk EVs inhibit αCD3/αCD28 induced proliferation of CD4+CD45RA+ T cells. Purified CD45RA+CD45RO-CD4 T cells of 2 different donors were labeled with CFSE and stimulated with 1.5 µg/ml αCD3 and 1 µg/ml αCD28 in the presence or absence of milk EVs and EV-depleted controls for 6 days. Cells were subsequently harvested and CFSE dilution was determined by flow cytometry. Bars represent mean ± SD, each T cell donor was cultured with 3 different milk donors.

**Supplementary Fig. 5:**
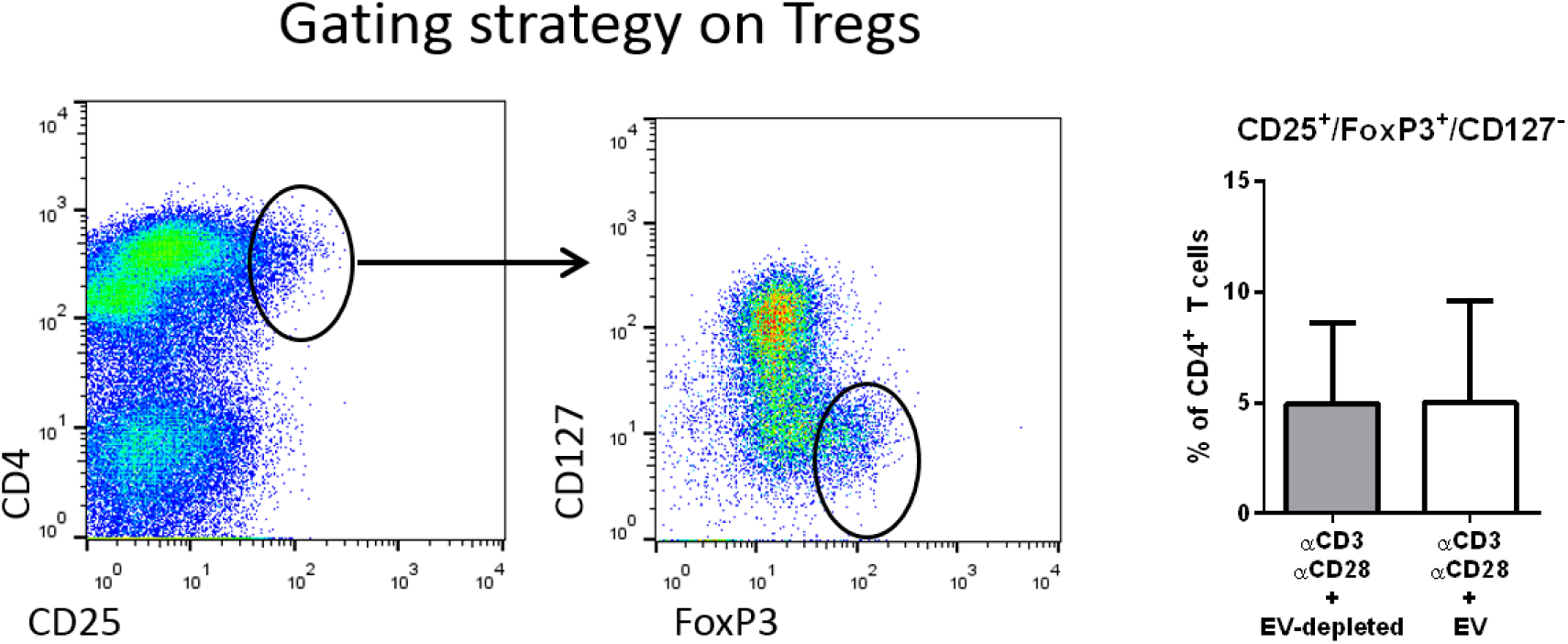
Human milk EVs do not induce a regulatory T cell phenotype. Quantification of the typical CD4+CD25highFoxP3+CD127-phenotype of human regulatory T cells in PBMC following exposure to 1.5 µg/ml αCD3 in the presence of EV or donor-matched EV-depleted controls. Bars represent mean ± SD of 4 independent experiments performed with 4 different PBMC donors and in total 6 different milk donors.

**Supplementary Fig. 6:**
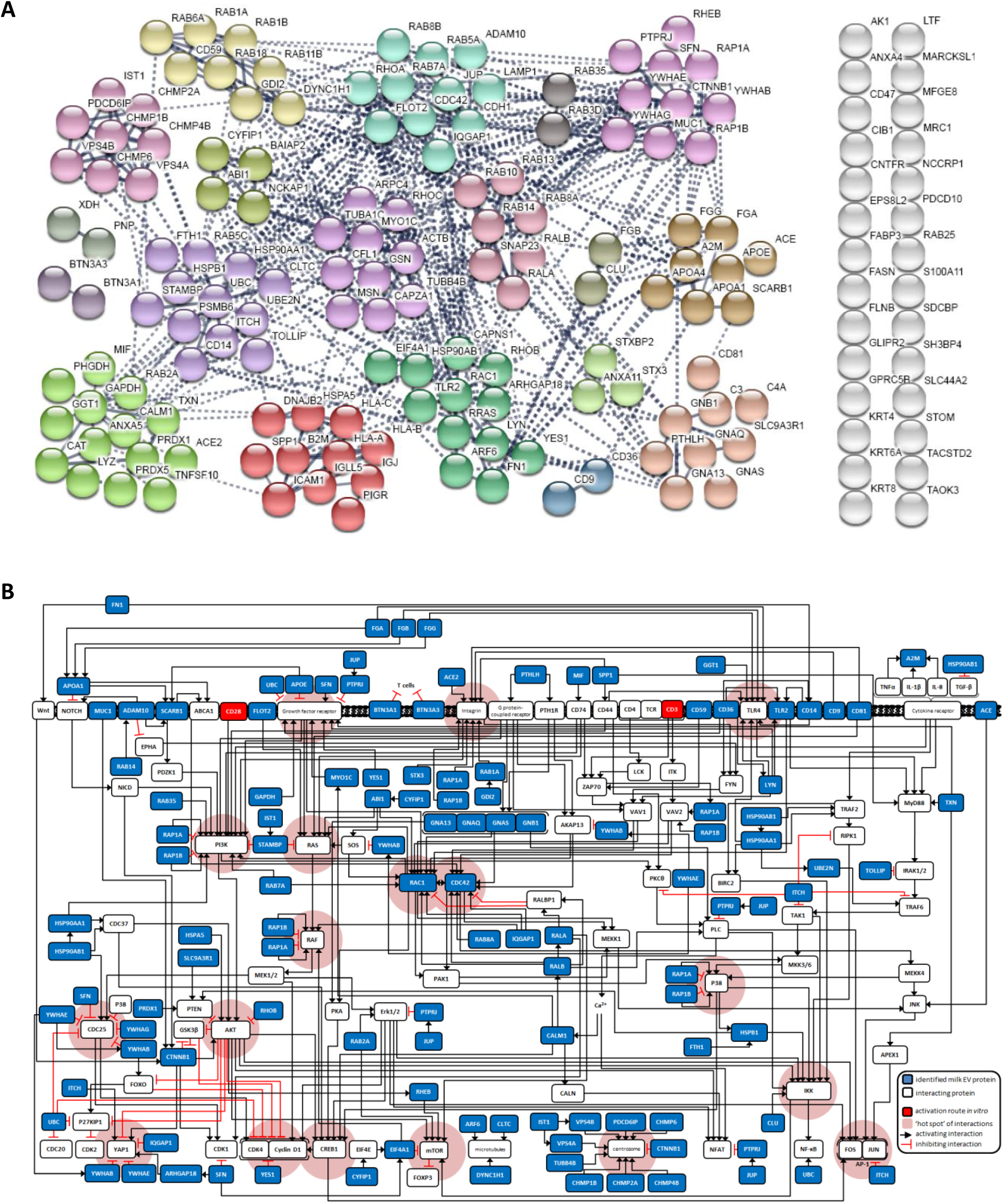
Enrichment and protein-protein network analysis allowed complete integration of human milk EV proteins into signaling pathways associated to regulation of T cell activation and revealed interactions at multiple levels in the signaling cascades. A) Protein-protein interaction analysis on the identified milk EV proteins that linked to the selected GO-terms was performed (minimum required interaction score set to high confidence 0.700), followed by k-means clustering (showing 19 clusters) in order to visualize the most likely occurring construct clusters within the network. A total of 137 proteins formed protein-protein interactions (which is 83.0%) and 28 proteins had no interaction with any other milk EV protein (which is 17.0%). Only those proteins that were part of an interaction network were further investigated for validated links to T cell activation. B) Functional annotation analysis of previously identified milk EV proteins that link to the observed *in vitro* effects in Fig. 3: inhibition of T cell activation. The interaction of selected milk EV proteins (in blue) and cellular proteins (in white; either shown with their common gene name, or a synonym when widely used in literature) and the type of interaction (activating or inhibiting) within relevant signaling pathways are shown. If a protein has interactions with ≥ 6 other proteins, this node is shown in red as a ‘hotspot’. Although milk EV proteins were selected via relevant GO-terms, some proteins could not be linked to the specific signaling cascades and are not shown in B (see Supplementary File 2 for a full overview of the analysis).

**Supplementary Fig. 7:**
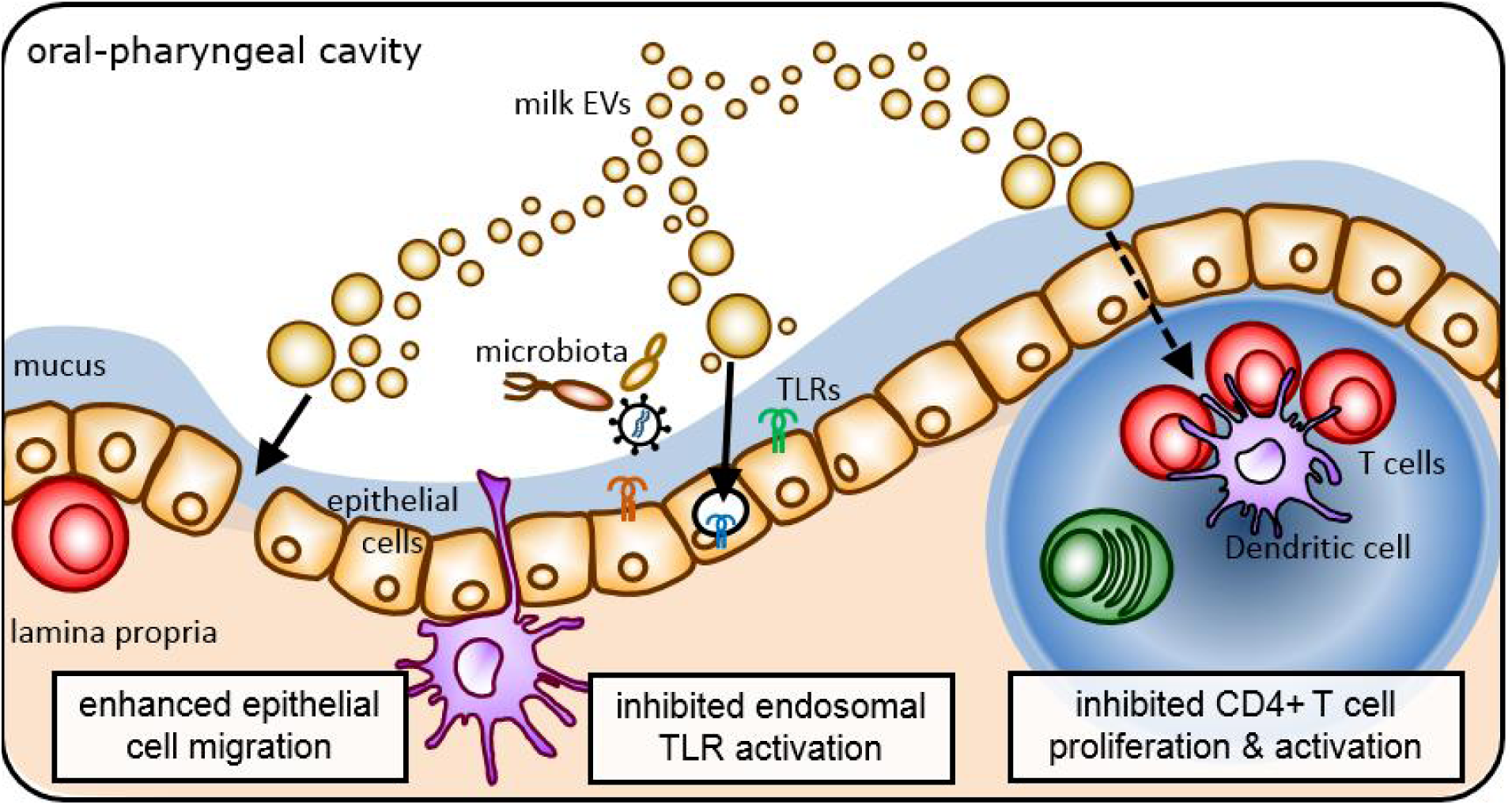
Graphical conclusion. Milk extracellular vesicles (EVs) are a heterogeneous population of cell-derived multi-component nanoparticles that can interact and modulate a variety of cells present in the infants oral mucosa. Together with the microbiota, possible pathogens, food- and environmental components, maternal milk EVs interconnectedly communicate with these mucosal cells. The gingival epithelial cells must form a tight barrier in order to prevent breaches in the mucosal surface. Milk EVs enhance the migratory capacity of epithelial cells, allowing for rapid gap closure. Additionally, epithelial cells are part of the innate immune system as they scan for microbe-associated molecular patterns (MAMPs) via Toll Like Receptors (TLRs). Milk EVs specifically tune TLR signaling by inhibiting endosomal TLR activation. Since EVs can cross epithelial barriers, they can also reach adaptive immune cells which reside underneath the epithelial layer. In the presence of milk EVs, CD4+ helper T cells are inhibited in their activation but remain responsive to stimulation once milk EVs are absent. The collective cargo of the heterogeneous population of maternal milk EVs works in concert to target key hotspots of signaling networks in target cells to specifically fine-tune mucosal processes involved in barrier function and immunity, thereby creating a window for adaptation and regulated development.

**Supplementary Fig. 8:**
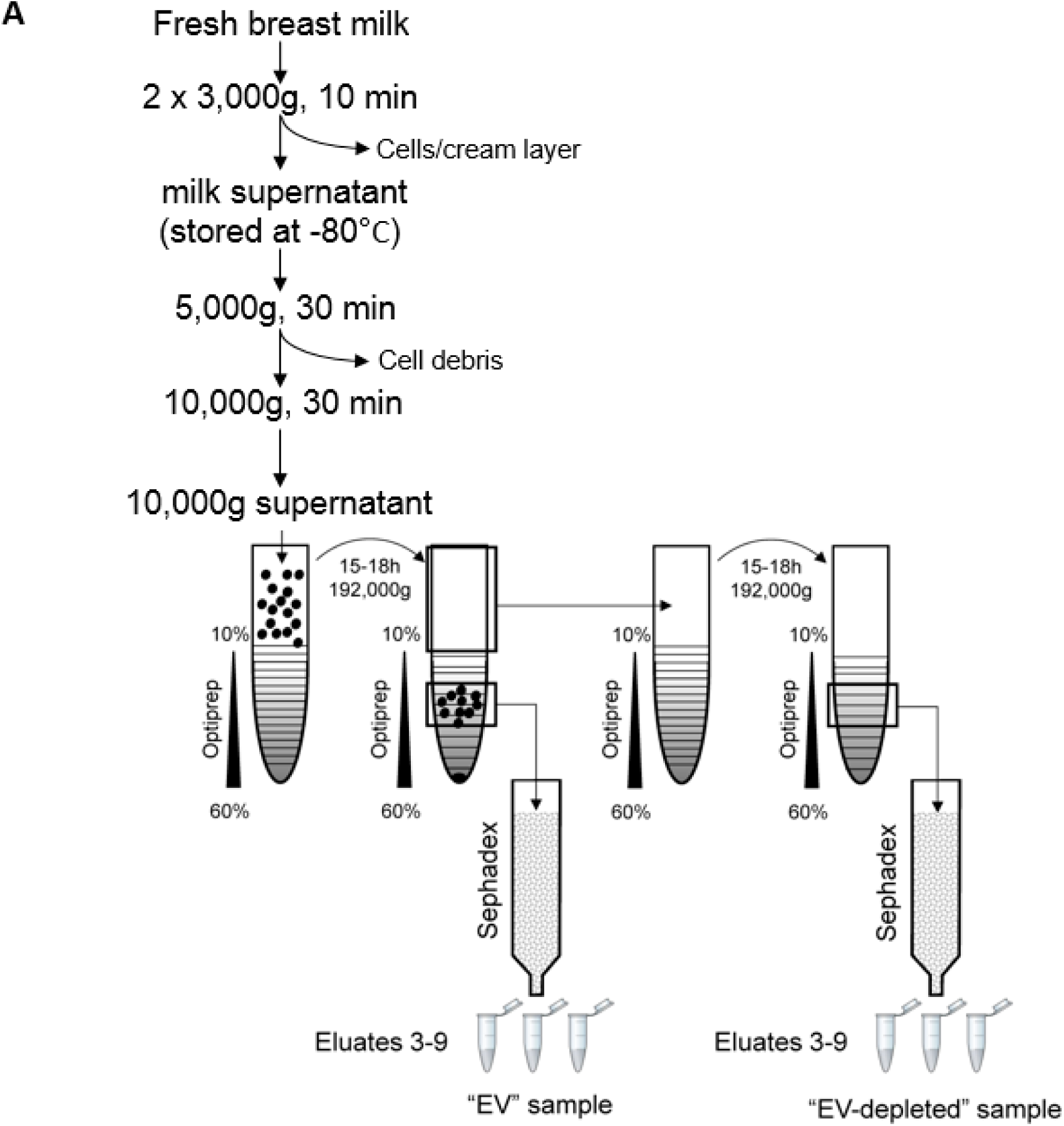
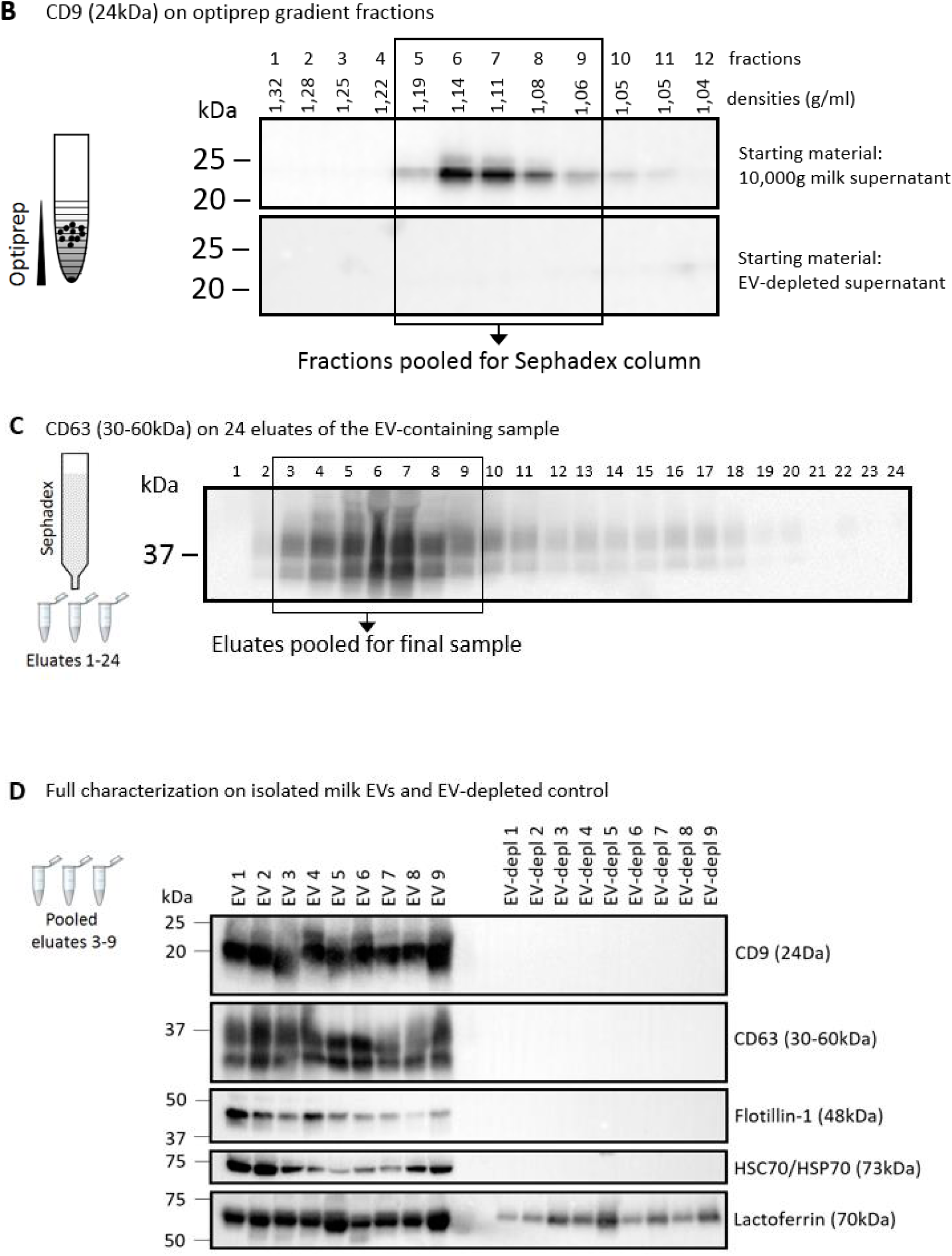
Isolation and characterization of human milk EVs and EV-depleted milk control. A) Schematic overview of the isolation of milk EVs and the EV-depleted milk control using differential centrifugation followed by density gradient ultracentrifugation. The original isolation procedure^7^ applied sucrose as a density medium. However, to maintain functionality of isolated EVs, Optiprep was used to build the gradient for ultracentrifugation. Size exclusion chromatography (SEC) using sephadex columns was applied to separate EVs from Optiprep and to collect the ‘EV fractions’ in the culture medium needed for functional analysis. B) Western blot analysis for CD9 (exposure time 60 seconds), together with buoyant density determination, was used to identify which fractions contained EVs. These fractions were pooled and introduced to the Sephadex column. C) Western blot analysis for CD63 (exposure time 20 seconds) and CD9 (data not shown) was used to determine which eluates contained EVs. D) The pooled eluates that were used as EV or EV-depleted samples were characterized by Western blot for the presence of EV-associated markers CD9 (exposure time 1 second), CD63 (exposure time 5 seconds), flotillin-1 (exposure time 120 seconds), Hsp70 (exposure time 30 seconds) and non EV-associated marker lactoferrin (exposure time 10 seconds) (data from 9 individual milk donors). The concentration of EVs used in the *in vitro* assays were within the physiological range since the starting volume of milk 10,000g supernatant was 6.5 ml prior to density gradient centrifugation and the volume of the pooled SEC eluates 3-9 was 7 ml. The volume of isolated milk EVs used in each *in vitro* assay was as high as possible.

## Acknowledgements

This work was supported by project 11676 within the framework of a partnership program jointly funded by Nutricia Research and the Dutch Technology Foundation STW, which is part of the Netherlands Organization for Scientific Research (NWO), and is partly funded by the Ministry of Economic Affairs (to M.I.Z.; M.J.C.H.; F.A.R.; E.N.M.N.; M.H.M.W.). The European Union’s Horizon 2020 Framework Programme under the grant FETOPEN-801367 evFOUNDRY (to M.J.C.H.; M.K.; M.H.M.W.). The European Union’s Horizon 2020 research and innovation Programme under the grant Marie Skłodowska-Curie-722148 TRAIN-EV (to A.G.; M.H.M.W.). The TiFN Food and Nutrition; project OH001, ‘novel strategies to promote oral health’ (to M.M.F.; M.K.). The European Research Council under the 441 European Union’s Seventh Framework Programme (FP/2007-2013)/ERC Grant Agreement number 442 337581 (to E.N.M.N.)

