## Supplementary material for "Human milk extracellular vesicles target nodes in interconnected signalling pathways that enhance oral epithelial barrier function and dampen immune responses": Suppl File 1

### Slide 1
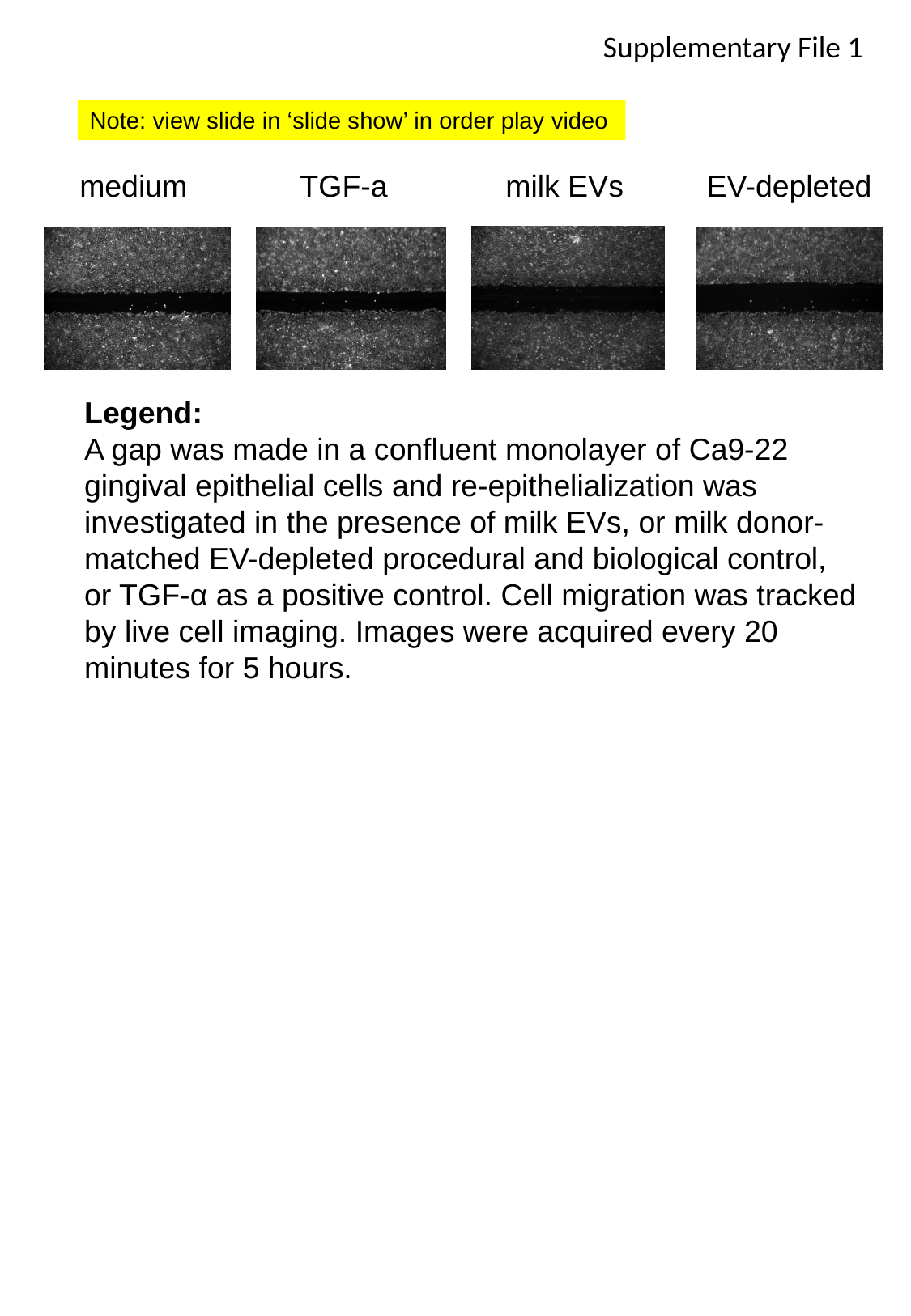

Supplementary File 1
Note: view slide in ‘slide show’ in order play video
medium
EV-depleted
TGF-a
milk EVs
Legend:
A gap was made in a confluent monolayer of Ca9-22 gingival epithelial cells and re-epithelialization was investigated in the presence of milk EVs, or milk donor-matched EV-depleted procedural and biological control, or TGF-α as a positive control. Cell migration was tracked by live cell imaging. Images were acquired every 20 minutes for 5 hours.
